# Seizures in a Bang-Sensitive Epilepsy Model Impact Associative Learning in *Drosophila* Larvae

**DOI:** 10.1101/2025.08.20.671221

**Authors:** Keller T. Gammons, Sej Cho, Elaine R. Reynolds

## Abstract

The seizures associated with epilepsy in humans may impact their learning and memory, however, the mechanism of how the seizures themselves lead to these deficits is not clear. *Drosophila* mutant flies that have seizure phenotypes have been used effectively as a model for human epilepsy. In this study, we tested whether stimuli that induce seizures in *Drosophila para*^*bss*^ mutants, a mutation in the fly sodium channel gene, affect learning. We used an adult conditioned courtship model and both rewarding and aversive associative learning assays in *Drosophila* larvae. For the conditioned courtship model, males exposed to mated females show depressed mating when placed with virgin females, suggesting that they learned from their rejection. In the larval paradigm, larvae are trained by pairing an odor (octanol or amyl acetate) with a reward (fructose) or an aversive stimulus (electrical shock) and then tested for learning by looking for movement towards or away from the odor they were trained on in a choice assay. We found that our wildtype CS stock and *para*^*bss*^ mutants did not respond to octanol as a neutral odor but as an aversive one. Therefore, octanol was used to train in a positive associate learning task and amyl acetate was used in aversive assays. Our *para*^*bss*^ stock performed almost as well as wildtype in learning in all of these assays, although other labs had reported learning deficits in these mutants. In addition, seizure induced through vortex in the adult learning paradigm and cold in the larval assay disrupted learning. These results suggest that learning deficits may be directly due to seizures rather than some other effect of the mutant.

**Author Summary:** In humans, seizures like those seen in epilepsy are associated with deficits in learning and memory. However, the link between the seizures and these deficits are not well understood and there is conflicting studies about what the characteristics of the seizure and the types of learning or memory affected. Animal model systems have been effectively used to study human disease and for this study, we chose the fruit fly, Drosophila melanogaster. *Drosophila* mutant flies that have seizures have been used to study human epilepsy and have similar types of learning and memory found in higher animals. We asked whether seizure induced in a fly mutant would disrupt learning and memory. We used several types of learning tasks, and both fly adults and larva. While the mutant flies performed almost as well as normal flies in all of these learning tasks, when they had seizures right after they learned a task, the learning was disrupted. These results suggest that learning deficits in humans may be directly due to seizures rather than something else

## Introduction

Seizures and related disorders such as epilepsy in humans are correlated with deficits in learning and memory and are associated with learning disabilities and other neuropsychiatric comorbidities [1–4]. Long term exposure to seizure can produce neurodegeneration. For example, the incidence of seizure has been correlated with cognitive decline in the development of Alzheimer’s disease [5–6]. In animal models, repeated brief seizures induce progressive hippocampal neuron loss and memory deficits [7]. Hyperexcitability also has been linked directly to Aβ42 expression in a fly Alzheimer’s Disease (AD) model suggesting a potential mechanism for the relationship [8]. Seizure mutants in flies have also been shown to have neurodegeneration and shortened lifespan, although the causal nature of that relationship has been questioned [9–10].

However, in humans there is some conflicting evidence on whether seizures directly impact learning and memory. In a human study where seizures were monitored, subjects who experienced seizures during the experimental period performed equally well on memory tasks as subjects who did not have seizures [11]. However, application of electroconvulsive shock therapy (ECT) has been shown repeatedly to disrupt short term memory and to create lasting changes in synaptic plasticity in both humans and animals [12–14]. Several studies in rats showed that a seizure prior to learning can impact the ability to learn but doesn’t affect longer term memory, although alternative timing or intensity of the seizure can impact memory [15–16]. In addition, Nygaard et al (2015) showed in a mouse model of epilepsy that an anti-epileptic drug improved learning and memory in mice models by regulation of the pattern of oscillations in brain circuits between excitatory and inhibitory activity [17]. Also, deep brain stimulation can improve cognition when used for seizure control [4]. So, there is much evidence for a link between seizure and cognition even if the nature of that relationship is not understood.

As a means of developing a model to look at causal or mechanistic connections between seizure and learning and memory processes, we used fruit fly models of seizure and learning. *Drosophila* has been used effectively as a model for human epilepsy by using seizure mutants that resemble genetic epilepsies seen in humans [18–19]. In this study, we tested whether seizures induced in *Drosophila para*^*bss*^ mutants affect learning. This bang-sensitive mutant has a mutation in the voltage-gated Na+ channel gene *para (paralytic)*, which lowers its threshold for seizures as compared to wildtype [20–21]. This sensitivity causes the flies to seize and paralyze in response to a mechanical stimulation or when exposed to the cold [22–23].

To assess learning we used an adult courtship learning assay and a larval olfactory learning assay. Both are associative memory tasks, defined as the ability to develop a relationship between unrelated stimuli or items. Adult courtship assay is a commonly used method in flies of assessing learning and memory [24–25]. In this assay, a negative association is generated. Males are exposed to mated females and therefore, they “learn” not to attempt mating since they have been repeatedly rejected by the females. In the courtship assay with a virgin female, the time to mate (mating depression) is extended by this exposure or males may fail to mate at all.

Another common way to study associative learning in *Drosophila* is the olfactory paradigm, which pairs a reward or aversive stimulus with an odor [26–27]. Larval associative learning assays using olfaction have been more recently pioneered [28–29]. Apostolopoulou et al (2013) showed that larvae can form associations between odors and appetitive gustatory reinforcement, like fructose [30]. Pauls et al (2010) found that larvae were able to associate olfactory stimuli with an electric shock punishment [31]. In our larval olfactory assays, we used both rewarding and aversive associations.

Our study looked for a causal relationship between seizure and learning disruption using both the adult and larval learning assays. We expected that seizure would disrupt learning in both models. In the adult assay, *para*^*bss*^ males were conditioned by the mated females and then seized by mechanical exposure before examination in the courtship assay. In the larval assay, *para*^*bss*^ larvae were trained by association of an odor with fructose or an electrical shock, exposed to the cold, and then their olfactory preference was examined. Seizures disrupted learning in both the adult and larval assays. This seizure learning model in flies can be used to understand the parameters of the disrupted learning and memory, and the underlying mechanisms.

## Methods

### Fly strains and maintenance

Wild-type Canton-S (CS), *dunce (dnc)*, and *paralytic*^*bss*^ (*para*^*bss*^) flies were raised on standard molasses/cornmeal food and kept on a 12:12 light cycle, at 25°C. The *para*^*bss*^ flies had been previously backcrossed and maintained in the CS background.

### Adult Learning Assay

#### Courtship paradigm

Male flies were moved into behavior chambers via weak suction and allowed to acclimate to the environment for 10 minutes. A virgin female fly was then introduced, also via suction, and the timer was started. Observations of two courtship measures were recorded in this study, time to chasing, defined in this study as continuous following for at least five seconds, and wing extension, as indicated by the 90° wing extension of the male that is used to produce a mating song. Copulation was noted if it occurred within the observational time frame. Behavior was recorded via a Leica EZ4W video microscope capture using Leica imaging software with a hard stop time set at 20 minutes and then timepoints were coded in a spreadsheet for all assays. Males that did not exhibit the measured behaviors in the allowed 20 minutes were recorded as having a time of 20 minutes in data processing [32].

#### Experimental design

Four conditions were tested using the CS and *para*^*bss*^ strain with 15 flies per condition. The first condition was just a standard courtship assay as described above. In the second condition, flies were exposed to mated females. Under these conditions, males learned not to mate as indicated by delayed courtship times. Males were placed into a standard vial with 10 previously mated females for 12-14 hours overnight and then tested for learning in the courtship assay. In the third condition, flies were seized by applying a mechanical shock for 10 seconds at high speed on a vortex mixer [19]. Flies were allowed to rest for 15 min after vortexing before courtship testing. Finally, in the fourth condition, flies were exposed to mated females, then vortexed as described above before the courtship assay was performed.

### Larval Learning Assay

#### Collection of Drosophila larva

Collection of larvae was done in accordance with the methods from Apostolopoulou et al (2013) and Pauls et al (2010) [30–31]. Larvae were collected 5-6 days post hatching once they reached their 3rd-instar feeding stage directly from the food. Larvae were washed and kept moist to avoid inducing pupation. Larvae were also tested to make sure they were moving before using in the learning paradigm.

#### Initial odor response

An initial odor preference was recorded for the strains of larvae used in the experiment. Initial odor preference was assessed twice. Standard 85 mm in diameter Petri dishes with 2.5% agar in PBS were used in the assay. Either amyl acetate (AM) or 3-octanol (OCT) was loaded into 4.5 mm diameter Teflon containers with perforated lids and placed onto one side of an agar plate. A group of 30 crawling 3rd-instar larvae were placed on the agar plate at the center of plate. Larvae were then given 5 min with the odor and were able to freely move around the agar plate. Larvae present on the half of the plate with the odor were counted. If greater than 60% were on the half with the odor, the strain was viewed as having a preference for the odor. 40-60% was viewed as neutral and less than 40% was viewed as aversive.

#### Choice Assays

Choice assays were used to determine both untrained and trained odor responses in CS and *para*^*bss*^. Odor choice was assessed twice. Standard 85 mm in diameter Petri dishes with 2.5% agar in PBS were used in the assay. AM or OCT were loaded into the Teflon containers with perforated lids. The agarose plate was divided into three sections, with the AM container on one side and the OCT container on the other side with the middle region as a neutral zone. Thirty crawling 3rd instar larvae were placed onto the neutral zone, and the petri plate lid was replaced to allow an olfactory gradient to develop. Larvae position was documented before and after the 5-minute trial time.

#### Rewarding Associative Learning

OCT was paired with a sugar reward. CS larvae were accessed in six trials, *para*^*bss*^ larva were accessed in seven trials. Standard petri plates for training were 2.5% agar in PBS with 2 M fructose in the plates. A group of 30 larvae were placed on the fructose agar plate with a Teflon container holding OCT. Larvae are left to explore the odor in the presence of the fructose reward for 5 minutes. They were then moved to a moist agar resting plate with no odor for 5 minutes. This process was repeated two more times. Once training was complete for a total of 3 times, 15 of the still moving larvae underwent a choice assay as described above. AM was also tested in a positive associative assay but a change in the choice assay was not observed and so all further experiments were performed with OCT. Trials were also conducted with AM and a rewarding stimulus (CS 6 trials, *para*^*bss*^ 7 trials).

#### Aversive Associative Learning

[AM](http://learning.am/) was paired with an aversive stimulus. CS larvae were accessed in four trials, *para*^*bss*^ larva were accessed in four trials. Consistent with Pauls et al. (2010), 85 mm in diameter Petri dishes were filled with 2.5% agar in PBS. Two semicircular copper electrodes (1 mm in diameter, 70 mm in length) were adjusted to the dimension of the Petri dish and immersed in the agar. The electrodes were arranged opposite to each other in the plate with a distance of 5 cm at their ends and 7.5 cm in the middle. Plates were then allowed to harden. AM was loaded into the Teflon containers with perforated lids. After a 4-minute acclimation period, larvae received a repeating shock of 60 V lasting 1 second for a total of 10/11 shocks in a 30 second period. Then larvae were transferred to a resting agar plate for a rest period for 5 minutes. The training process was then repeated two additional times. After the aversive odor to shock association training has been completed, larvae were given an additional 5-minute resting period. The 15 selected larvae that displayed normal mobility were then tested in the choice assay. OCT was also tested in a positive associative assay but a change in the choice assay was not observed and so all further experiments were performed with AM. Trials were also conducted with OCT and an aversive stimulus (CS 4 trials, *para*^*bss*^ 4 trials).

#### Production of Seizures

To test the effect of seizure on the larval learning process, 15 selected larvae were placed on an empty plate on top of ice for five minutes before the choice assay. With this procedure seizures are observed in the *para*^*bss*^ strain but not the CS. The larvae were then given five minutes on the rest plate to allow them time to return to their normal temperature and mobility before the choice assay. In the OCT rewarding assay, CS larvae were accessed in six trials, *para*^*bss*^ larvae were accessed in seven trials. In the AM rewarding assay, CS larvae were accessed in six trials, *para*^*bss*^ larvae accessed in seven trials. In the AM aversive assay, CS larvae were accessed in three trials, *para*^*bss*^ larvae accessed in three trials. In the OCT aversive assay, CS larvae were accessed in three trials, *para*^*bss*^ larvae accessed in three trials.

#### Data Collection

Images at the beginning and the end of each choice assay were taken on iPhone 13Pro live photo, and the position of the larvae were counted and recorded in a spreadsheet. Larvae were counted in the 3 zones on the agar plate: left zone (AM), middle zone (Neutral), and right zone (OCT). The percent preference for each trial was calculated and then averaged across trials. For each strain of larva, three conditions were examined: untrained choice preference, trained choice preference, and trained +cold choice preference.

#### Data analysis

All statistical analyses were conducted using RStudio (2022.12.0). In the adult courtship learning assay, means and standard deviations were calculated for time to wing extension for each of the groups. Findings were analyzed using a nonparametric two-way t test between groups using a p-value threshold of ≤ 0.05 for significance. In the choice assays, means and standard deviations were calculated for larval odor preference for each group. An ANOVA was conducted to examine the effects of strain (CS vs. *para*^*bss*^) and training condition (untrained, trained, trained with cold-induced seizure) using ≤ 0.05 as a threshold for significance. Post hoc analysis was completed using nonparametric two-way t test between groups using a p-value threshold of ≤ 0.05 for significance.

#### Data Availability Statement

All the data used for analysis in this paper is available as a supplement entitled S1Dataset.

## Results

### Adult Learning Assays

The adult learning assay results showing time to wing extension are shown in Figure 1. The *para*^*bss*^ flies learned successfully (condition 1 vs. condition 2, *p<0*.*0001*) and seizure was found not to affect the standard courtship assay (condition 1 vs, condition 3, *p=*0.611). There was a significant effect of seizure on learning (condition 2 vs condition 4, *p=*0.01). *Para*^*bss*^ flies mated a bit slower than CS in the standard courtship assay, although not significantly (CS 75 sec vs. 143 sec, p=0.75). Time to following data was collected and although it was somewhat more variable, it showed the same trends.

**Figure 1.**
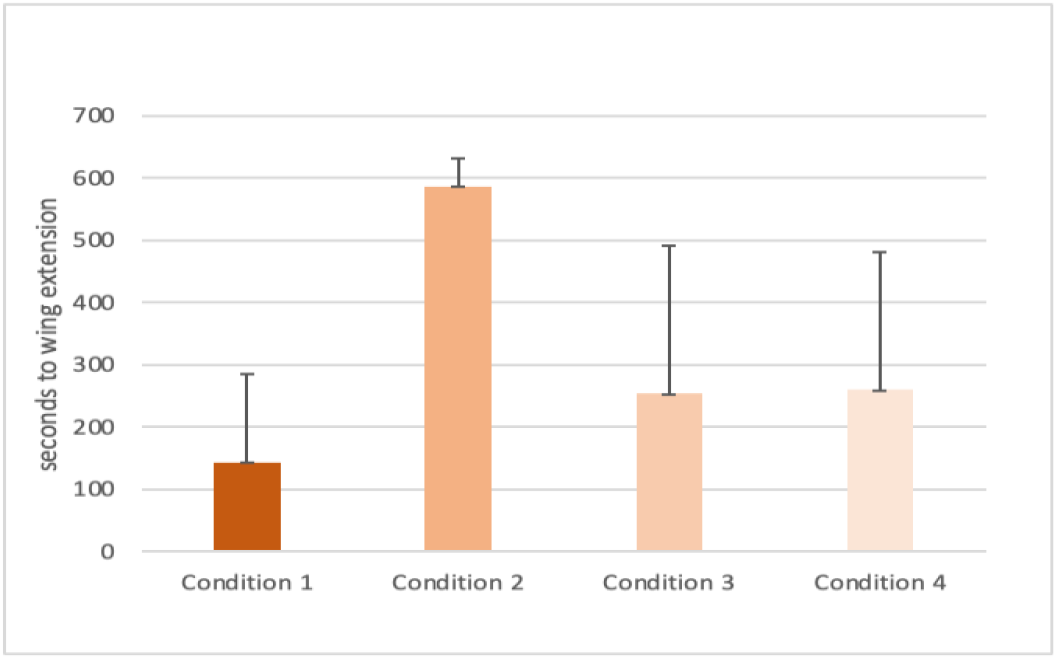
Adult courtship depression in the *para*^*bss*^ mutant using time to wing extension measure. Condition 1 Standard Courtship assay, Condition 2 Courtship depression after exposure to a mated female (p<0.0001), Condition 3 Standard courtship after exposure to cold, Condition 4 Courtship depression with exposure to mated female is not observed after exposure to cold (p=0.01).

### Larval Learning Assays

#### Preference Testing

We tested odor preference in the wildtype and mutant larvae in preparation for the learning assay (Figure 2). In non-competition testing with a single odor, CS wildtype and *para*^*bss*^ larvae displayed neutrality or a slight preference for the AM odor (CS AM preference, 60% **+** 4.7, and *para*^*bss*^ AM preference 53% **+**0) consistent with reports in the literature [4]. However, in contrast to this reference, CS and *para*^*bss*^ showed aversive behavior towards OCT (13% **+** 9.4, preference for *para*^*bss*^ and 37% **+** 4.7 for CS).

**Figure 2.**
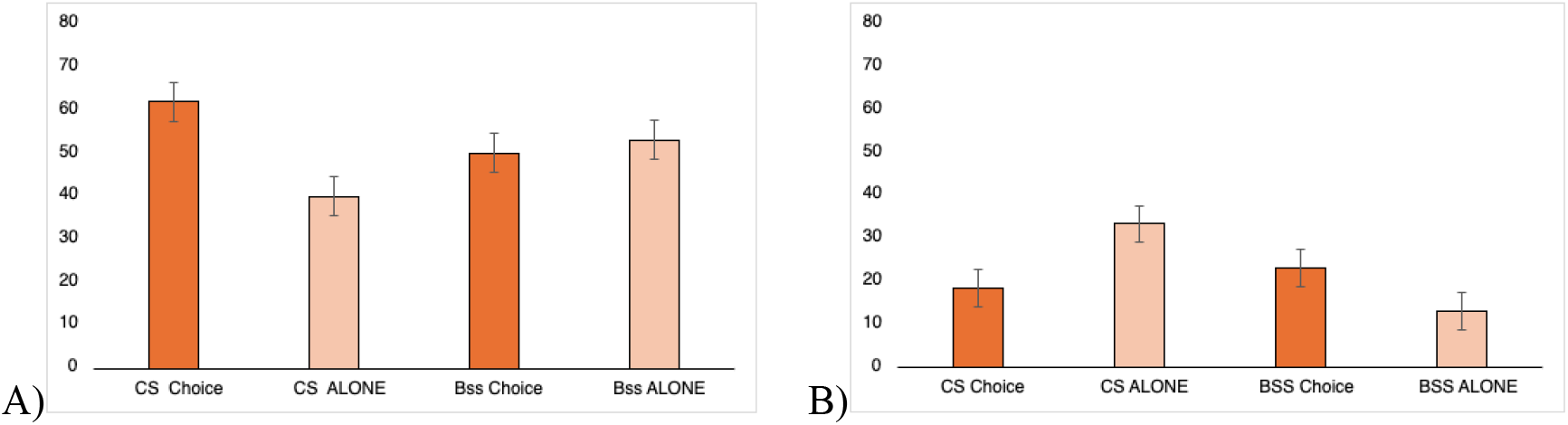
Initial odor preference for CS and *para*^*bss*^ strains. (A) Untrained preference for the amyl acetate odor alone or in a choice assay, (B) The untrained preference for the 3-octanol odor alone or in a choice assay. Error bars represent standard deviation.

In an untrained competition or choice assay, CS wildtype and *para*^*bss*^ larvae displayed a slight preference for the AM odor in the competition preference testing with OCT with percent preference of 62%**+** 10.4 and 50%**+** 0 respectively. On the OCT side of the plate, only 18%**+** 7.6 of CS and 23%**+** 5.8 of *para*^*bss*^ larvae were found, confirming the aversive nature of OCT in our assay. The remaining larvae were found in the neutral zone. The untrained choice assay for both the CS wildtype and *para*^*bss*^ larvae were used as the baseline initial preferences for the larvae in the positive and negative learning assays. Our original plan was to use *dunce (dnc)* as a negative learning control. However, upon doing initial preference testing, *dnc* mutant larvae did not display neutrality toward either odor (AM 29% **+** 4.0, and OCT 29%**+** 4.0). Since the strain’s behavior was quite different from our CS or *para*^*bss*^ strains, we decided not to include *dnc* results in our analysis.

#### para^bss^ can learn in both positive and negative assays

The *para*^*bss*^ larva demonstrated learning in both the rewarding and aversive associative olfactory assays (Figure 3). The *para*^*bss*^ larvae showed learning in the rewarding associative assay with OCT odor, displaying statistically significant increases in the choice assay post rewarding training from 23%**+** 5.8 to 59%**+** 8.6, (p= 0.047). The learning in *para*^*bss*^ larvae was comparable to that in the wildtype for OCT (18.3%**+** 7 to 55**+** 12.6). Learning was not demonstrated with either strain in the reward assay with AM, possibly because of a ceiling effect caused by our assay conditions.

**Figure 3.**
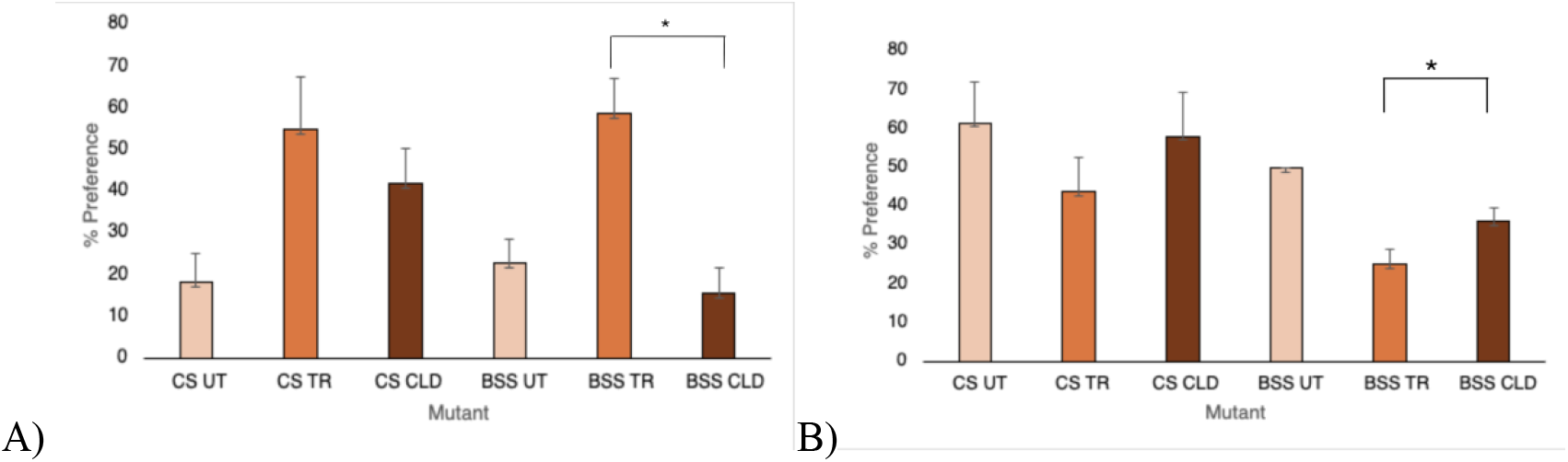
Rewarding and aversive larval learning assays with and without seizure A) Rewarding assay, (UT) no OCT training, (TR) trained with OCT paired with fructose, and (CLD) trained with OCT paired with fructose and then exposed to cold plate before choice assay. B) Aversive assay, (UT) no AM training, (TR) trained with AM paired to shock, and (CLD) trained with AM paired to shock and exposed to cold plate before choice assay. Error bars indicate the standard deviation. p<0.5 are represented by the ^*^ in post hoc comparisons.

The *para*^*bss*^ larvae also demonstrated learning in the aversive assays using the AM odor. In pairing the odor with an aversive stimulus, *para*^*bss*^ larvae demonstrated a significant decrease in percent preference for the AM odor results. Percent preference decreased from 50%**+** 0 to 25%**+** 4.1 post aversive training (p = 0.041). CS wildtype larvae demonstrated a comparable decrease in percent preference from 62%**+** 10.4 initially to 44%**+**8.9. Aversive training did not significantly reduce preference for OCT for either *para*^*bss*^ or CS.

#### Cold disrupts learning in para^bss^ in both positive and negative assays

Exposing *para*^*bss*^ flies to a cold shock, which induces visible seizures, disrupts learning in both the rewarding and aversive associative learning assays. In the rewarding assay, there was no main effect of strain (p = 0.097) and a significant main effect of training condition (p = 0.004). However, post hoc analysis showed a significant impact of cold on *para*^*bss*^ learning (p = 0.019) but not on CS learning (p = 0.7). In the aversive assay, there was a significant main effect of strain (p = 0.002) and a significant main effect of training condition (p = 0.002). Post hoc analysis showed a significant impact of cold on *para*^*bss*^ learning (p = 0.03) but not on CS learning (p = 0.07). The post hoc analysis confirms that cold-induced seizures in *para*^*bss*^ larvae significantly impact learning in the choice assay.

## Discussion

We were able to demonstrate learning in *para*^*bss*^ mutants using both a traditional adult courtship learning assay and in larval learning assays. In the courtship assay, the *para*^*bss*^ flies show the expected latency to mate, indicating that they were able to learn the negative association. In addition, *para*^*bss*^ larvae also were able to learn in associative olfactory learning assays using aversive and rewarding stimuli. In the rewarding setup, pairing the OCT odor with a fructose solution during training generated *para*^*bss*^ larval learning, demonstrated by an increased preference for the OCT odor. In the aversive assays, the pairing of an aversive shock treatment with the AM odor also generated associative olfactory learning, with *para*^*bss*^ showed an average decrease in preference for the AM odor.

In previous publications, *para*^*bss*^ as well as a knockout of the *paralytic* gene showed decreased ability to learn in an aversive phototaxic suppression assay [33–34]. Quinine was used as the aversive odor, and the flies were 10 days old. In experiments with our strains, *para*^*bss*^ did not perform well on phototaxic assays, moving towards light at a lower rate than control flies in preliminary control experiments. In addition, both our CS and *para*^*bss*^ strains showed neutrality to quinine. So, it was not possible for us to replicate these previous studies. One possible difference that may account for the alternative results in adult flies is that our experiments were performed with newly emerged flies aged 5 days, while previous studies used flies that were 10 days old. Since the adult seizure phenotype worsens with age in these mutants, this may lead to a learning deficit with age. In addition, later learning deficits might be due to neuronal degeneration and we wanted to focus on the impact of the seizure on learning in this study. The larval paradigm then alleviates variability that might be observed with aging adult flies.

Induced seizure in the *para*^*bss*^ mutant showed a disruption of both adult and larval learning. In the adult courtship paradigm, seizure induced by vortex reduced the time to initiate courtship to levels observed in the control courtship assay, alleviating the courtship depression induced by learning. In the larval assays where seizure was induced by exposure to cold, training no longer produced a change in preference for both associative learning paradigms. Preference for odors in both the positive and aversive assay returned to pretraining levels in the *para*^*bss*^, however exposure to the cold did not disrupt learning in the CS wildtype control.

While the larval learning assay is promising in allowing exploration of simple associative learning, we saw some limitations for its use. Previous research had labeled the odors used in our experiments as neutral [30]. We found that OCT was repulsive in both individual and choice assays. Therefore, we could not use OCT in our aversive assay, and the results are presented only using AM. AM was closer to neutral when tested alone, but in a choice assay paired with OCT it was consistently the preference. In rewarding assays with AM, we seemed to have a ceiling effect, where training did not improve AM preference in the choice assay. Therefore, our rewarding assays were done with OCT. These changes in preference from published work could be due to strain differences or limitations of the assay conditions. Our CS and *para*^*bss*^ strains have roughly the same background since *para*^*bss*^ was isolated in the CS background and was also backcrossed to maintain the CS background as a control in behavioral experiments. We also originally sought to use *dunce (dnc)*, a mutant that doesn’t display learning as a negative control in these experiments. However, *dnc* shows different preferences in both single odor and choice assays from our CS and *para*^*bss*^ strains. It is possible that in addition to learning deficits, *dnc* might have defects in odor perception.

The model developed in our study of *Drosophila* can be used to understand and to discover the basic mechanisms that link seizure and learning. Drosophila has historically contributed to an understanding of the molecular pathways that underlie learning and memory [28]. More recently, Sears and Broadie (2025) have used the Drosophila excitability mutants to look at mechanisms behind learning in Kenyon cell circuits, suggesting that excitability’s impact on learning may be due to interactions between PKA and ERK signaling pathways [35]. Though the differences in the human and the invertebrate fly model are significant, the fly model allows for detailed investigations that separate different variables in this complex relationship.

